# Reactomics: using mass spectrometry as a reaction detector

**DOI:** 10.1101/855148

**Authors:** Miao Yu, Lauren Petrick

## Abstract

Untargeted metabolomics analysis captures chemical reactions among small molecules. Common mass spectrometry-based metabolomics workflows first identify the small molecules significantly associated with the outcome of interest, then begin exploring their biochemical relationships to understand biological fate (environmental studies) or biological impact (physiological response). We suggest an alternative by which biochemical relationships can be directly retrieved through untargeted high-resolution paired mass distance (PMD) analysis without a priori knowledge of the identities of participating compounds. Retrieval is done using high resolution mass spectrometry as a chemical reaction detector, where PMDs calculated from the mass spectrometry data are linked to biochemical reactions obtained via data mining of small molecule and reaction databases, i.e. ‘Reactomics’. We demonstrate applications of reactomics including PMD network analysis, source appointment of unknown compounds, and biomarker reaction discovery as a complement to compound discovery analyses used in traditional untargeted workflows. An R implementation of reactomics analysis and the reaction/PMD databases is available as the pmd package (https://yufree.github.io/pmd/).

## Introduction

Untargeted metabolomics or non-targeted analysis using high resolution mass spectrometry (HRMS) is one of the most popular analysis methods for unbiased measurement of organic compounds ^1,2^. A typical metabolomics sample analysis workflow will follow a detection, annotation, MS/MS validation and/or standards validation process, from which interpretation of the relationships between these annotated or identified compounds can then be linked to biological pathways or disease development, for example. However, difficulty annotating or identifying unknown compounds always limits the interpretation of findings ^3^. One practical solution to this is matching experimentally obtained fragment ions to a mass spectral database ^4^, but many compounds remain unreported/absent, thereby preventing annotation. Rules or data mining-based prediction of in silico fragment ions is successful in many applications ^2,5^, but these approaches are prone to overfitting the known compounds, leading to false positives. Ultimately such workflows require final validation with commercially available or synthetically generated analytical standards, which may not be available, for unequivocal identification.

Potential molecular structures could be discerned using biochemical knowledge, through the integration of known relationships between biochemical reactions (e.g. pathway analysis) ^3^. Such methods are readily used to annotate compounds by chemical class. For example, the Referenced Kendrick Mass Defect (RKMD) was able to predict lipid class using specific mass distances for lipids and heteroatoms ^6^, and isotope patterns in combination with specific mass distances characteristic of halogenated compounds such as +Cl/-H, +Br/-H were used to screen halogenated chemical compounds in environmental samples ^7^. For these examples, known relationships among compounds were used to annotate unknown compounds, as a complementary approach to obtaining compound identifications.

The most common relationships among compounds are chemical reactions. Substrate-product pairs in a reaction form by exchanging functional groups or atoms. Almost all organic compounds originate from biochemical processes, such as carbon fixation ^8,9^. Like base pairing in DNA ^10^, organic compounds follow biochemical reaction rules, resulting in characteristic mass differences between the paired substrates and their products. We present a new concept, paired mass distance (PMD), that reflects such reaction rules by calculating the mass differences between two compounds or charged ions. PMD can be used to extract biological inference without identifying unknown compounds.

Exploiting mass differences for compound identification is not new. Mass distances have been used to reveal isotopologue information when peaks show a PMD of 1 Da ^11^, adducts from a single compound ^12^ such as PMD 22.98 Da between adducts [M+Na]^+^ and [M+H]^+^, or adducts formed via complex in-source reactions ^13^ from mass spectrometry data. Such between-compound information has also been used to make annotations of unknown compounds ^4,14^, to classify compounds ^15^ or to perform pathway-independent metabolomic network analysis ^16^. However, these calculations of PMD were used to identify compounds or pathways in ultimately facilitate interpretations of the relationships between important compounds. Here, we propose that PMD can be used directly, skipping the step for annotation or identification of individual compounds, to aggregate information at the reaction level, called ‘Reactomics’.

HRMS can directly measure PMDs with the mass accuracy needed to provide reaction level specificity. Therefore, HRMS has the potential to be used as a reaction detector to enable reaction level study investigations. Here, we use multiple databases and experimental data to provide a proof-of-concept for using mass spectrometry in ‘reactomics’. We also discuss potential applications such as PMD network analysis, biomarker reaction discovery, and source appointment of unknown compounds. We envision that these applications will reveal the measurable reaction level changes without the need to assign molecular structure to unknown compounds.

## Results and Discussion

### Definitions

We first define a reaction PMD (PMD_R_) using a theoretic framework. Then we demonstrate how a PMD_R_ can be calculated using KEGG reaction R00025 as an example (see Equation 1). There are three KEGG reaction classes (RC00126, RC02541, and RC02759) associated with this reaction, which is catalyzed by enzyme 1.13.12.16.

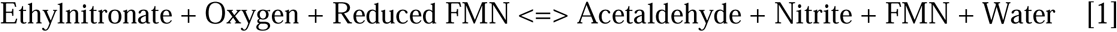

In general, we define a chemical reaction (PMD_R_) as follows:

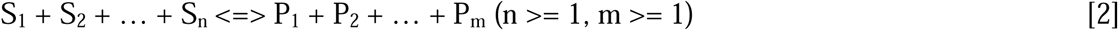

Where: S means substrates and P mean products, and n and m the number of substrates and products, respectively. A PMD matrix [M1] for this reaction is generated:

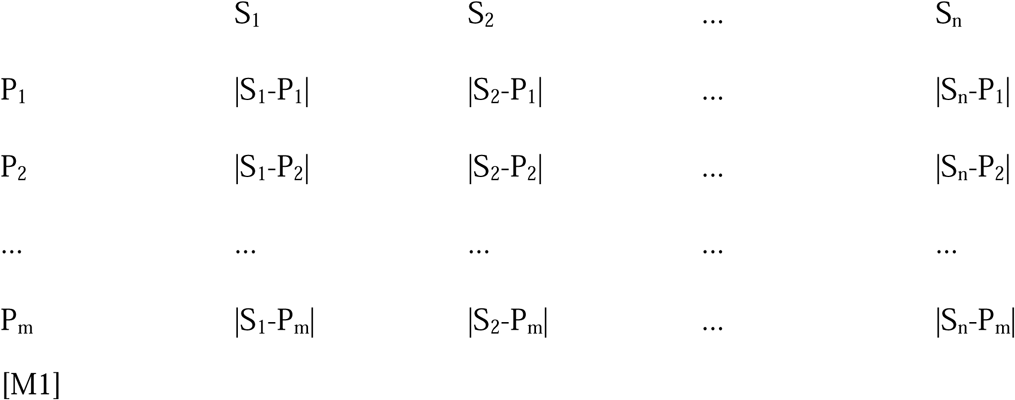

For each substrate, S_k_, and each product, P_i_, we calculate a PMD (|S_n_-P_m_|).

Assuming that the minimum PMD would have a similar structure or molecular framework between substrate and products, we select the minimum numeric PMD for each substrate as the substrate PMD (PMD_Sk_) of the reaction (Eq. 2).

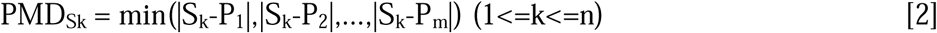

Then, the PMD_R_, or overall Reaction PMD, is defined as the set of substrates’ PMD(s) (Eq. 3):

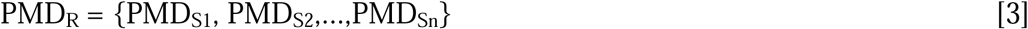

For KEGG reaction R00025, S_1_ is Ethylnitronate, S_2_ is Oxygen, S_3_ is Reduced FMN, P_1_ is Acetylaldehyde, P_2_ is Nitrite, P_3_ is FMN, P_4_ is Water, n=4, and m =3. A PMD matrix [M2] for this reaction can be seen below, where we define PMD_Ethylnitronate_ = 27.023 Da, PMD_Oxygen_ = 12.036 Da and PMD_Reduced FMN_ = 2.016 Da.

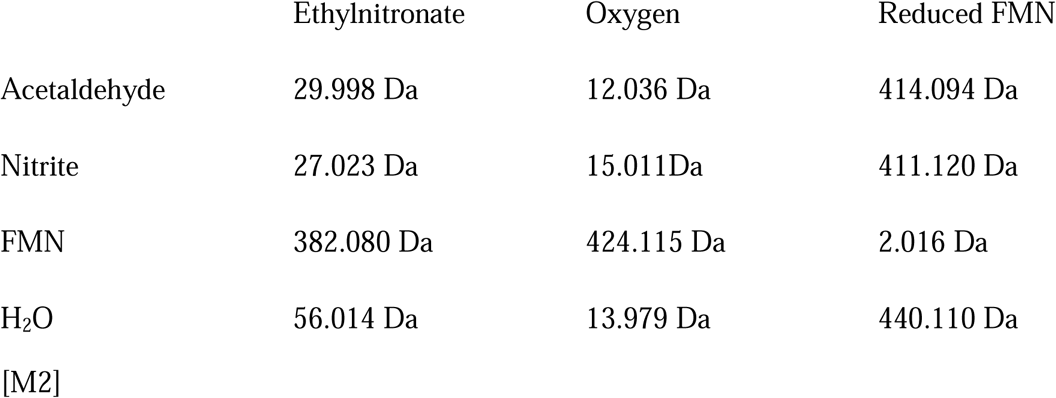

In our example, there are three PMD_R_ calculated from three PMD_S_: PMD_R_ is 27.023 Da, which is equivalent to the mass difference between two carbon atoms and three hydrogen atoms: PMD_R_ is 12.036 Da for the additions of two carbon atoms and four hydrogen atoms and loss of one oxygen atom: and PMD_R_ is 2.016 Da for the addition of two hydrogen atoms. However, other reactions may have multiple PMD_S_ that generate the same PMD_R_ value, such as certain combination reactions or replacement reactions. In this case, only one value will be kept as reaction PMD. In addition, each PMD_R_ has two notations. One is shown as an absolute mass difference of the substrate-product pairs’ exact masses or monoisotopic masses with unit Da. Another notation is using elemental compositions as the differences between two chemical formulas. Here, we describe it as an elemental composition instead of chemical formula, because it also describes the gain and loss of elements, and therefore the neat mass change. In our example reaction, the PMD_R_ can also be written as +2C3H, +2C4H/-O, and +2H, respectively. This elemental composition can be linked to known chemical processes retrieved from a reaction database, ie. KEGG. For example, +2H represents the elemental composition change of a reaction involving a double bond breaking such as KEGG example RC00126, and +2C3H indicates reaction with nitronate monooxygenase (EC:1.13.12.16) or reaction class RC02541. However, some elemental compositions, such as +2C4H/-H in our example, might not have a clear mechanism (e.g. no suggested KEGG reaction selection). By this definition, PMD_R_ can be generated automatically in terms of elemental compositions or mass units in Da.

We used these definitions to establish reference databases of PMDs. We used KEGG as a ‘reaction database’representing common reactions in human endogenous pathways, and we used human metabolome database (HMDB)^17^ as the ‘compound database’ representing common reactions between chemicals measured in human biofluids (see S.1 in the supplementary information).

### Qualitative and quantitative PMD analysis with mass spectrometry

PMD can be determined in biological or environmental samples from peaks observed in mass spectrometry. Mathematically, a PMD of uncharged compounds is equivalent to the PMD of their charged species observed with a mass spectrometer, as long as both compounds share the same adducts, neutral losses, and charges. In example reaction [1], reduced FMN has a monoisotopic mass of 458.1203 Da, while FMN has a monoisotopic mass of 456.1046 Da. Spectra from HMDB^17^ showed that common ions for reduced FMN and FMN using liquid chromatography (LC)-HRMS in negative mode are typically [M-H]^−^ with m/z 457.1124 and 455.0968, respectively. The mass distance of the monoisotopic masses is 2.016 Da and the mass distance of the observed adducts is also 2.016 Da. In cases such as this, mass spectrometry can be used to detect the PMD of paired compounds, but only for HRMS (see S.1 for data mining, S.2 for issues dealing with redundant peaks and fragments, and S.3 for discussion of resolution issues).

In addition to qualitative analysis, peaks that share the same PMD can be summed and used as a quantitative group measure of that specific ‘reaction’ in the sample, thereby providing a description of chemical reaction level changes across samples without annotating individual compounds. We define two types of PMD across samples: static PMD in which intensity ratios between the pairs are stable across samples, and dynamic PMD in which the intensity ratios between pairs change across samples. Only static PMDs, those with similar instrument response, can be used for quantitative analysis (see Table S.4 for theoretical example). Similar to other non-targeted analysis^18^, an RSD between quantitative pair ratios < 30% and a high correlation between the paired peaks’ intensity (> 0.6) are suggested to be considered a static PMD.

### PMD network analysis

Using the proposed PMD network analysis (see S.4 for details), we can identify metabolites associated with a known biomarker of interest. In fact, PMD network analysis can also be used in combination with classic identification techniques to enhance associated networks with targeted biomarkers. As a proof of concept, we re-analyzed data from a published study to detect the biological metabolites of exposure to Tetrabromobisphenol A (TBBPA) in pumpkin ^19,20^ using a local, recursive search strategy (see Figure 1). Using TBBPA as a target of interest, we searched for PMDs linked with the debromination process, glycosylation, malonylation, methylation, and hydroxylation, which are phase II reactions (e.g. primary metabolites) found in the original paper. Using this PMD network analysis, we identified 22 unique m/z ions of potential TBBPA metabolites, confirmed by the presence of brominated isotopologue mass spectral patterns (Figure S1.). This total was 15 more than the 7 unique ions that were described in the original publication. Such a network was built based on the experimental data and our local fast recursive search algorithms as shown in S.4. We found that most of the potential metabolites of TBBPA were found as higher-generation TBBPA metabolites, which are too computationally intensive to be identified using *in silico* prediction and matching protocols ^21^.

**Figure 1.**
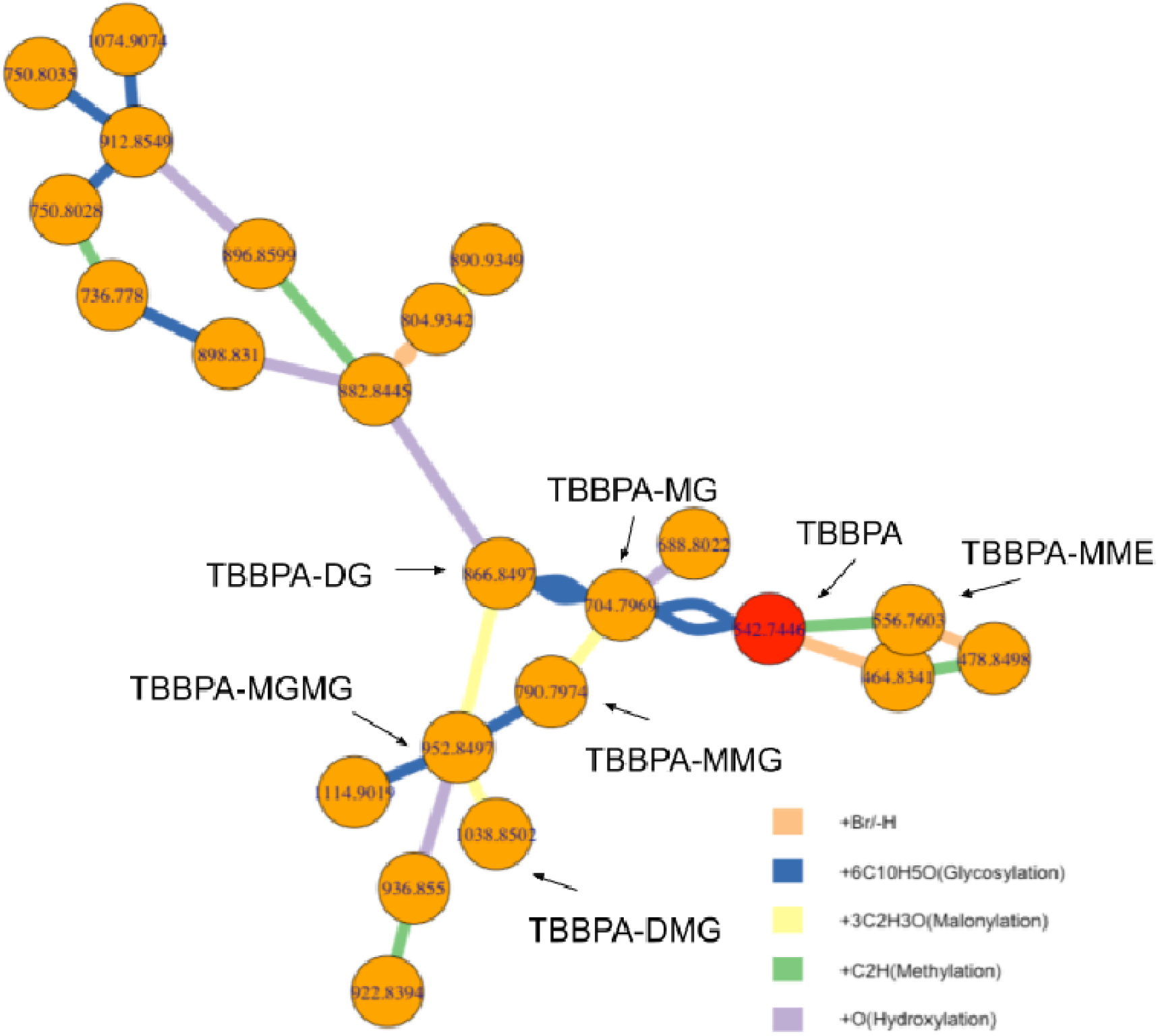
Metabolites of TBBPA in pumpkin seedlings’ root samples. Edges between two nodes were defined as Pearson’s correlation coefficients > 0.6 and shared reactions related PMDs including 77.91 Da for Debromination, 162.05 Da for Glycosylation, 86 Da for Malonylation, 14.02 Da for Methylation, and 15.99 Da for Hydroxylation. Previous reported metabolites are labeled.

### Source appointment of unknown compounds

When an unknown compound is identified as a potential biomarker, determining whether it is associated with endogenous biochemical pathways or exogenous exposures can provide important information toward identification. As can be seen in Tables S1 and S3, high frequency PMDs from HMDB and KEGG are dominated by reactions with carbon, hydrogen, and oxygen suggesting links to metabolism pathways. Therefore, if an unknown biomarker is mapped using a PMD network, connection to these high frequency PMDs would suggest an endogenous link. However, separation from this network is expected for an exogenous biomarker in which the reactive enzyme is not in the database, the exogenous compound is secreted in the parent form, or if it undergoes changes in functional groups such as during phase I and phase II xenobiotic metabolism processes. In this case, endogenous and exogenous compounds should be separated by their PMD network in samples.

Topological differences in PMD networks for endogenous and exogenous metabolites were explored using compounds from The Toxin-Toxin-Target Database (T3DB) ^22^. As shown in Figure 2, the PMD network of compounds was generated based on top ten high frequency PMDs of 255 endogenous compounds with 223 unique masses and 705 exogenous compounds with 394 unique masses and carcinogenic 1, 2A, or 2B classifications. Most endogenous compounds (Figure 2, red) were connected into a large network, while the exogenous compounds’ networks were much smaller (Figure 2, blue). Interestingly, most carcinogenic compounds were not connected by high frequency PMDs. Expanding this beyond just carcinogenic compounds, we randomly sampled 255 exogenous compounds from a total of 2491 exogenous compounds available in T3DB, and built a PMD network with the top 10 high frequency PMDs of those 510 compounds (255 exogenous compounds and 255 endogenous compounds). This step was repeated 1000 times, and the average degree of connection with other nodes was calculated as 4.5 (95% CI [4.3, 4.8]) for endogenous compounds and 1.7 (95% CI [1.2, 2.2]) for exogenous compounds.

**Figure 2.**
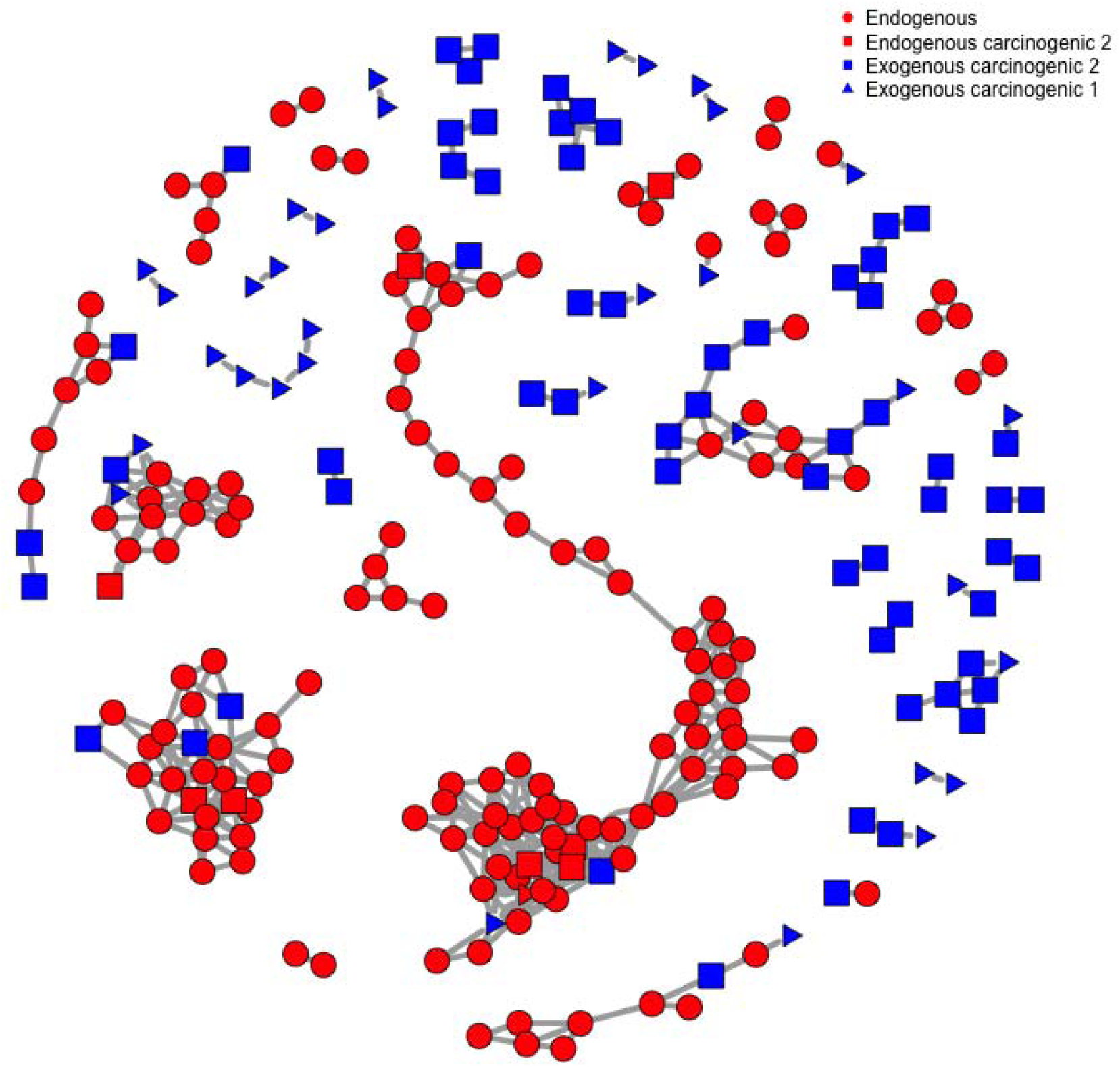
PMD network of 255 endogenous compounds and 705 exogenous carcinogens from T3DB database with the top 10 high frequency PMDs network. Compounds without linkage to other compounds have been removed.

Similar findings were observed for known compounds. We selected caffeine, glucose, bromophenol, and 5-cholestene as well characterized chemicals that are commonly observed with mass spectrometry, and paired them with other metabolites in the KEGG reaction database using the top ten high frequency PMDs from Table S1. As shown in Figure 3, different topological properties (e.g. number of nodes, average distances, degree, communities, etc.) of compounds’ PMD network were observed for each selected target metabolite. Endogenous compounds such as glucose or 5-cholestene were highly connected (average degree of node is 3.4 and 3.2, respectively) while exogenous compounds such as caffeine and bromophenol have more simple networks (average degree of node is 2.2 and 2.4, respectively). Further, the average PMD edge numbers between all nodes (edges end-to-end) in glucose and 5-cholestene networks are 9.7 and 6.6, respectively, while the average PMD edge numbers for caffeine and bromophenol are 3.3 and 1.8, respectively. Larger average PMD edge numbers mean a complex network structure with lots of nodes while smaller average PMD edge numbers mean a simple network structure with a few nodes. Based on these estimates, we proposed that unknown metabolites with average network node degree more than 3 would be likely endogenous compounds. Similarly, if the unknown compound belongs to a network with longer average PMD edge numbers, such compounds might also be of endogenous origin.

**Figure 3.**
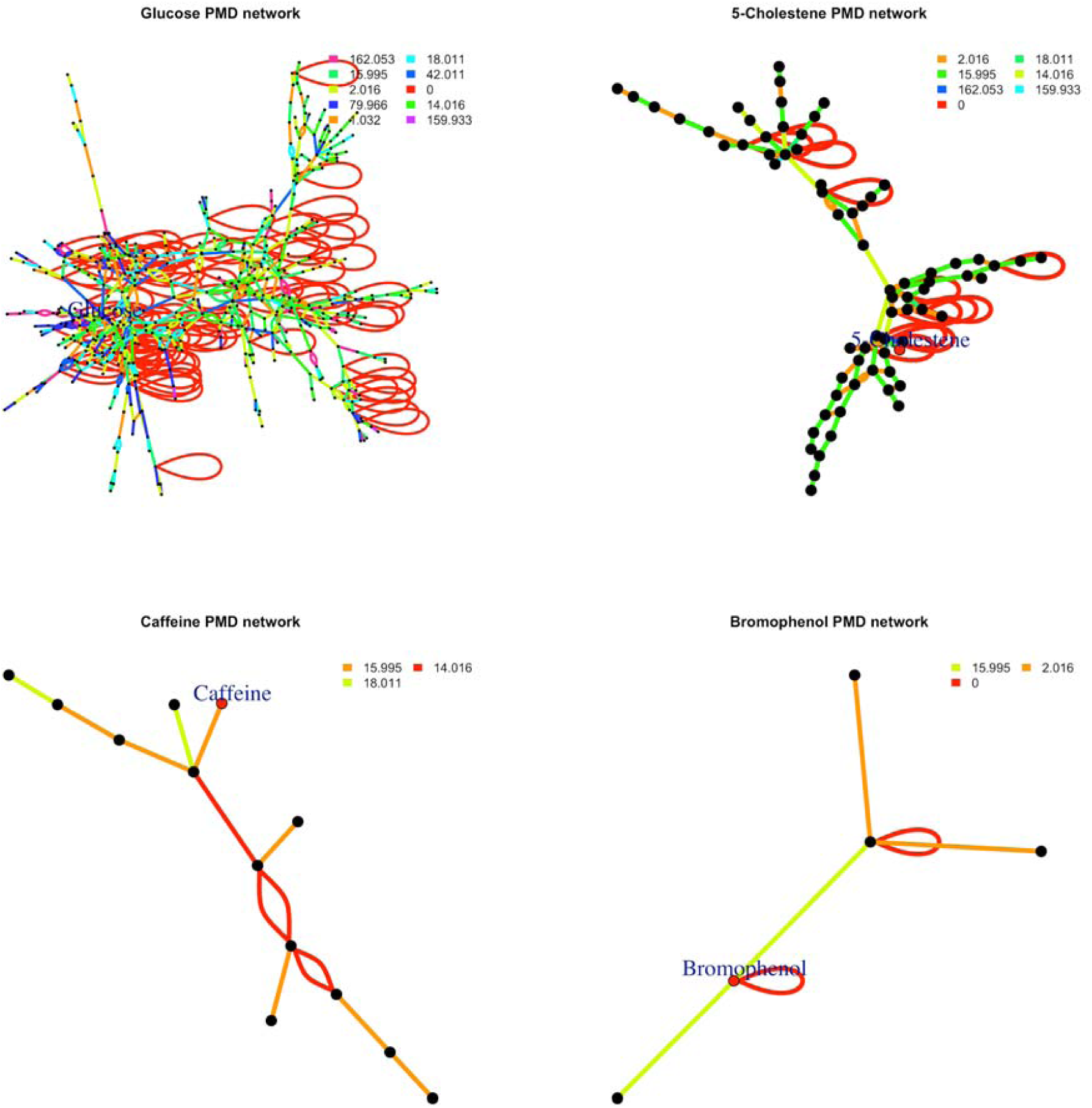
PMD networks for selected compounds from the KEGG reaction database. Networks are limited to relationships with the 10 top frequency PMDs.

### Biomarker reactions

Reactomics can be used to discover biomarker ‘reactions’ instead of biomarker ‘compounds’. Unlike typical biomarkers that are a specific chemical compound, biomarker reactions contain all peaks within a fixed PMD relationship and correlation cutoff. Thus, quantitative PMD analysis (see S.2) can be used to determine if there are differences between groups (e.g. control or treatment, exposed or not-exposed) on a reaction level. Such differences are described as a biomarker ‘reaction’.

We used publicly available metabolomics data (MetaboLight ID: MTBLS28) collected on urine from a study on lung cancer in adults^23^. Four peaks out of 1807 features from 1005 blood samples (469 cases and 536 controls) generated the quantitative responses of PMD 2.02 Da. This biomarker reaction (e.g. +2H from our annotated database) was significantly decreased in case samples compared with the control group (t-test, p < 0.05): see Figure 4). The original publication associated with this dataset did not report any molecular biomarker associated with this reaction ^23^, or the metabolites linked with this reaction, suggesting that quantitative PMD analysis offers additional information on biological differences between the groups on the reaction level that may be lost when focused on analysis at the chemical level. PMD-level investigations directly reduce the high dimensional analysis typically performed on a peaks or features level into low dimensional analysis on the chemical reaction level with explainable elemental compositions. Furthermore, these results suggest that follow-up analysis in this population should include targeted analysis of proteins or enzymes linked with +2H changes.

**Figure 4.**
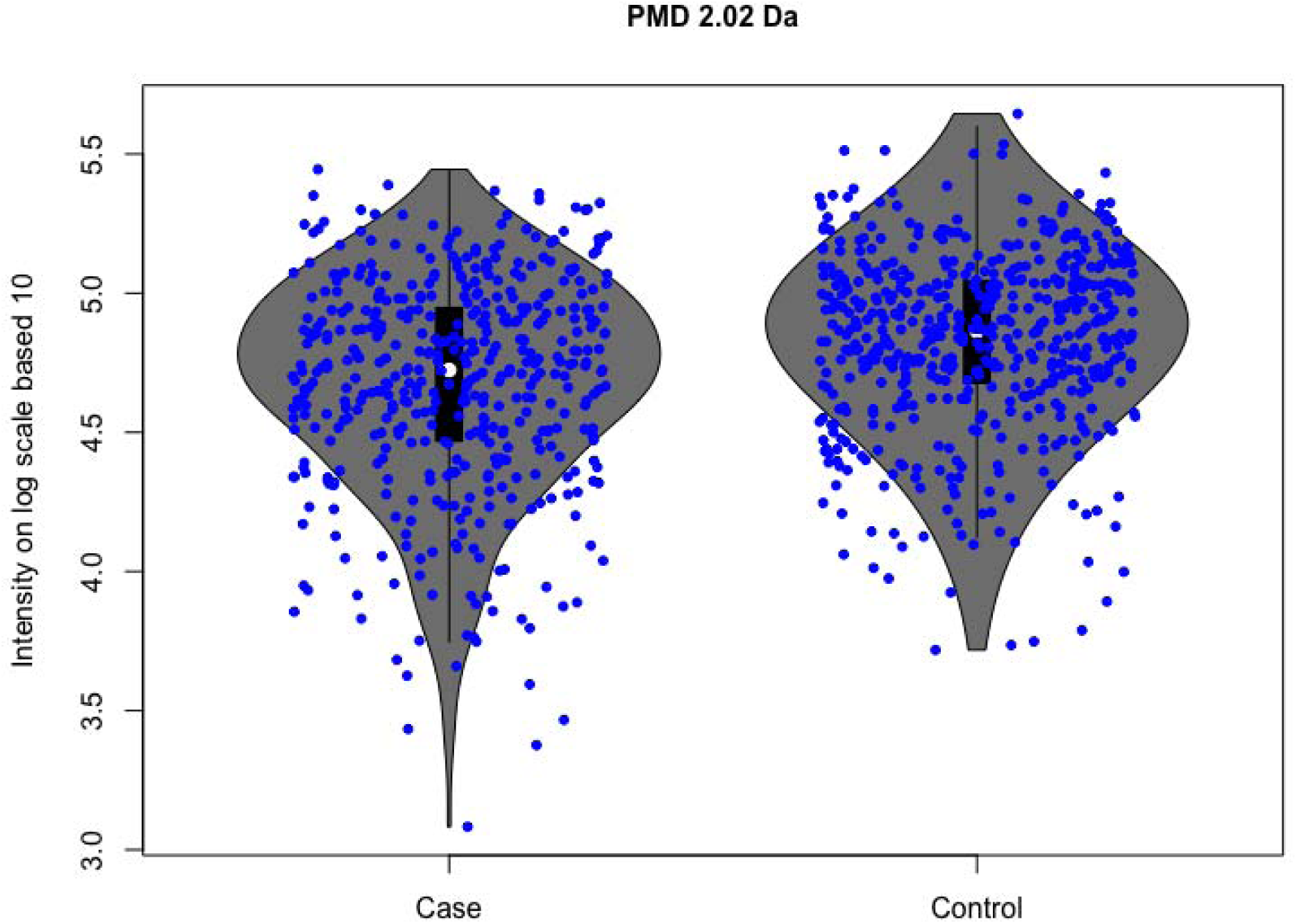
Quantitative PMD analysis identifies PMD 2.02 Da as a potential biomarker reaction for lung cancer (MTBLS28 dataset).

## Conclusion

We provide the theoretical basis and empirical evidence that high resolution mass spectrometry can be used as a reaction detector through calculation of high resolution paired mass distances and linkage to reaction databases such as KEGG. Reactomics, as a new concept in bioinformatics, can be used to find biomarker reactions or develop PMD networks. The major limitation of reactomics analysis is that mass spectrometry software is designed for analysis of compounds instead of reactions. In this case, the uncertainty in PMD measurements can not be captured directly from the instrument, and instead are calculated after data acquisition. Nevertheless, reactomics techniques provide information on biological changes for new biological inferences that may not be observed through classic chemical biomarker discovery strategies.

## Acknowledgments

This research was financially supported by NIEHS grants P30ES23515, 1U2CES030859, R21ES030882, and R01ES031117.

## TOC

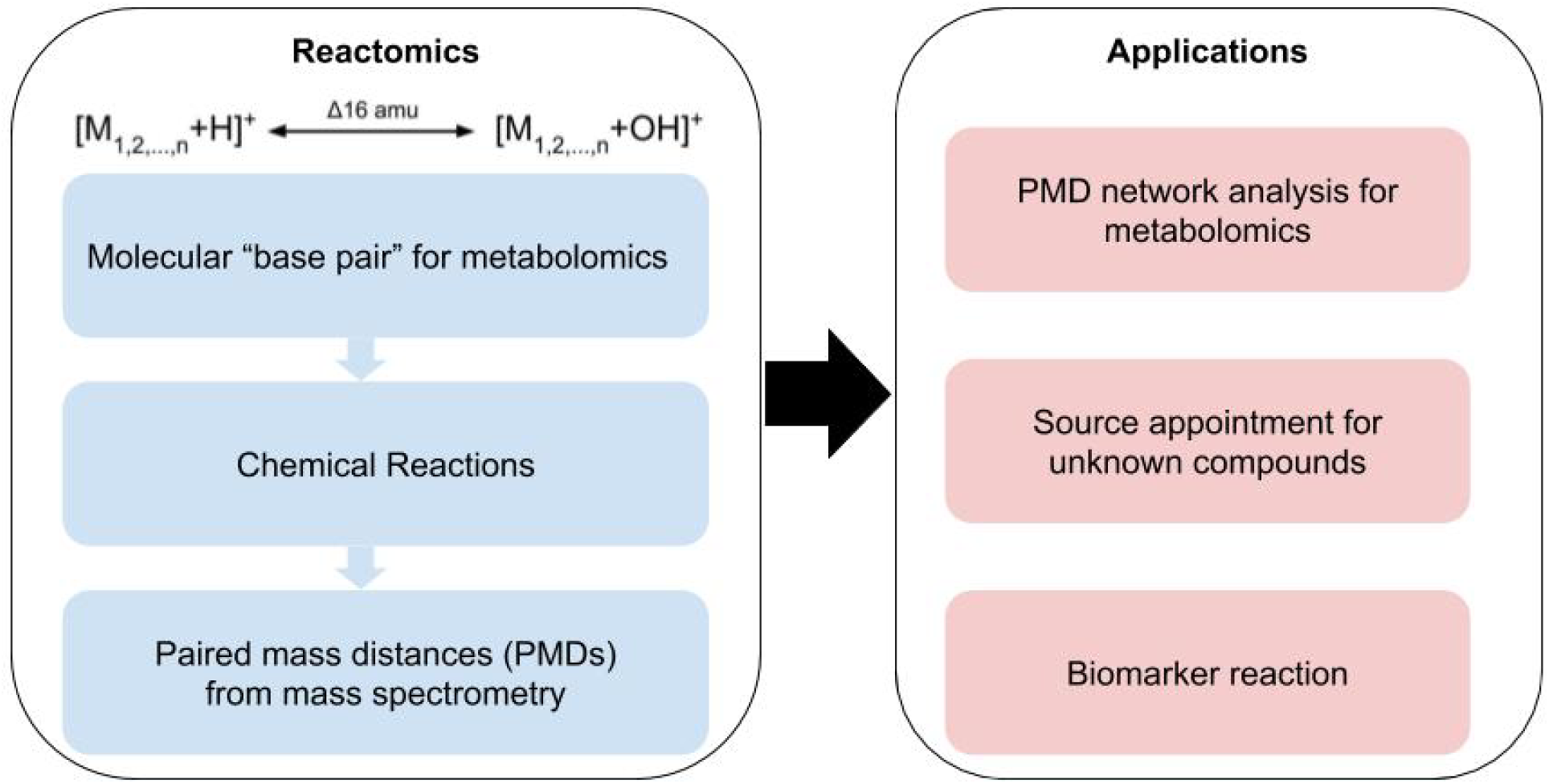

