## Supporting Information for "Reactomics: using mass spectrometry as a reaction detector"

**S.1 Data mining**

KEGG, with 11416 reactions, was used as a reference reaction database ^1^. All product and substrate molecules with exact mass larger than 100 Da and smaller than 1000 Da were retained, and PMDs associated with metals or ions of PMDs were removed. Then, we calculated PMD_R_ for the 9200 remaining KEGG reactions and identified 2548 unique reaction PMDs (in Da, reported to three decimal places). There are several common PMD_R_ values; the 10 highest frequency values covered 5448 KEGG reactions with frequency larger than 200. As shown in Table S1, high frequency PMDs were directly associated with similar biochemical reactions such as oxidation, breaking of double bonds, phosphate transfer reaction, etc. This unique property of PMDs facilitates annotation of reaction class or reaction-associated enzymes between a pair of compounds without a priori knowledge of the identity of each compound. Furthermore, PMDs of very low frequency can be used as biomarkers of unique reactions. Using the KEGG database as described, we generated a PMD database for reference annotation that is included in our open source software pmd package. Previous studies also use PMD calculated from the KEGG database for compound annotation ^2,3^. However, they did not formally define PMD_R_ , which would lead to spurious PMD calculations among ion pairs without clear biological or chemical meaning.

Compound databases including the human metabolome database (HMDB) ^4^ and The Toxin-Toxin-Target Database (T3DB) ^5,6^ were also employed to explore the PMDs among known compounds. There were 114100 compounds in HMDB and 3673 compounds in T3DB at the time of this data analysis. Unlike reaction databases, compound databases can reveal potential structure relationships or unknown reactions among compounds that may not be involved in specific endogenous human pathways. T3DB is the only database with annotation of their entries as endogenous or exogenous origin.

To demonstrate qualitative and quantitative PMD analysis, compounds from HMDB were used. First, the database was filtered to remove compounds with rare elements and those compounds not likely to be observed during mass spectrometry analysis. This resulted in 9516 unique formulas with carbon, hydrogen, oxygen, nitrogen, phosphorus and sulfur atoms and exact masses larger than 100 Da and smaller than 1000Da which were screened for PMD analysis.

**S.2 Redundant peaks and fragments in Reactomics**

A key issue in reactomics is redundant PMDs. Mass spectrometry collects peak- or feature-level data, many of which represent multiple signals from the same compound. Consequently, this will introduce redundant PMDs. For example, in positive mode, [M+H]^+^ will show a PMD 15.995 Da with [N+H]^+^ if M and N have an oxidation relationship. If we also collect the isotopologue of [M+H]^+^ and [N+H]^+^, their reaction level analysis would also indicate a PMD 15.995 Da leading to an overall false enrichment. Fortunately, these ions can be accounted for using the presence of PMD 14.995 Da and 16.995 Da as redundant PMDs. Similar scenarios will happen for adducts, neutral losses, and background peaks.

To avoid those redundant PMDs, annotation of the isotopologue,adducts, or other redundant peaks is needed for PMD analysis. This can be done using a variety of available softwares, including psuedu-spectra to annotate peaks based on known PMDs from annotation tools such as CAMERA ^7^, RAMclust ^8^, and InterpretMSSpectrum ^9^; or using the GlobalStd algorithm ^10^ or mz.unity ^11^ to annotate or remove redundant peaks based on PMD frequency analysis to capture unknown adducts. Using one of these algorithms, a single peak representing the same type (adduct, neutral loss, or isotope) between paired analytes can be selected for each cluster of redundant peaks. When the resulting filtered peaks are used for PMD analysis, they can then be linked to a specific biological reaction (PMD_R_) instead of redundant PMDs.

Another source of spurious PMDs is through relationships with fragment ions. Fragment ions can be generated during tandem mass spectral collection or through hard ionization processes. While fragmental patterns can be used to predict the structure of certain compounds with similar structures ^2,12,13^, reactomics, as proposed in this study, will not cover PMDs from fragment ions for compound identification. Here, reactomics is focused on PMD relationships among different compounds and their linkages to chemical reactions. In this case, full scan mode with soft ionization mass spectrometry data can only be used for analysis with reactomics, and MS data obtained using tandem mass spectrometry or hard ionization processes should be removed before the analysis of reaction level changes.

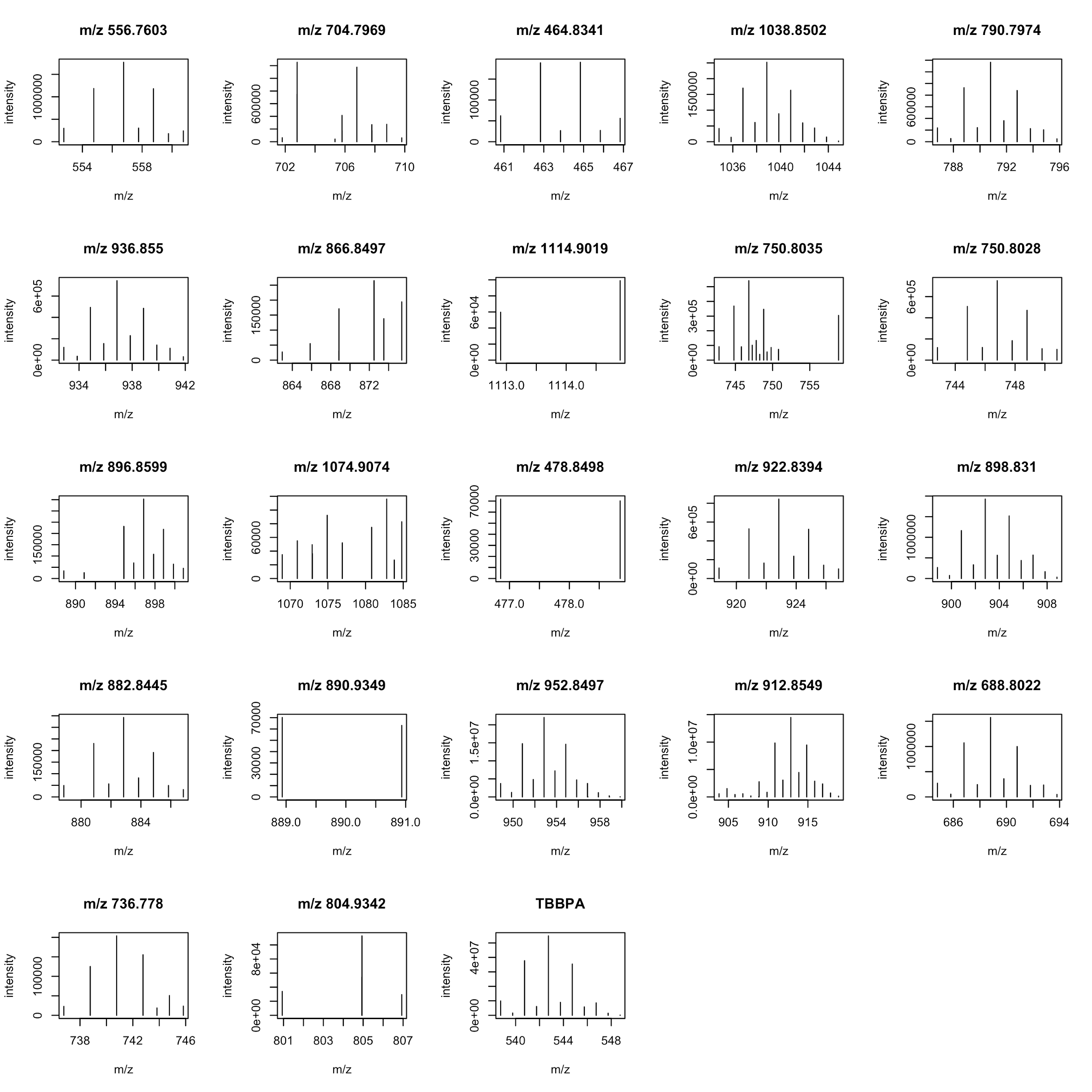

Figure S1. The peaks’ mass spectra with brominated isotopologues patterns from the pumpkin study.

Table S1. Top 10 high frequency KEGG reaction PMD_R_ and corresponding example reaction, reaction class, and enzyme.

| PMD (Da) | Freq | Example  Reaction class | Example  enzyme | Example reaction |
| --- | --- | --- | --- | --- |
| 2.016 | 1732 | RC00095 | 1.3.1.84 | NAD(+) + propanoyl-CoA <=> acryloyl-CoA + H(+) + NADH |
| 15.995 | 1169 | RC00046 | 1.3.99.18 | 2-Quinolinecarboxylic acid + Acceptor + H2O <=> 4-Hydroxy-2-quinolinecarboxylic acid + Reduced acceptor |
| 79.966 | 729 | RC00002 | 3.6.1.3 | ATP + H2O <=> ADP + H(+) + phosphate |
| 14.016 | 594 | RC00060 | 1.5.3.1 | S-Adenosyl-L-methionine + Glycine <=> S-Adenosyl-L-homocysteine + Sarcosine |
| 0 | 532 | RC00302 | 5.1.1.3 | L-glutamate <=> D-glutamate |
| 18.011 | 359 | RC00680 | 3.5.2.5 | (S)-Allantoin + H2O <=> Allantoate |
| 162.053 | 365 | RC00049 | 3.2.1.23 | H2O + lactose <=> D-galactose + D-glucose |
| 159.933 | 243 | RC02056 | 4.2.3.42 | 9alpha-Copalyl diphosphate + H2O <=> Aphidicolan-16beta-ol + Diphosphate |
| 1.032 | 262 | RC00006 | 1.4.1.2 | L-Glutamate + NAD+ + H2O <=> 2-Oxoglutarate + Ammonia + NADH + H+ |
| 42.011 | 237 | RC00004 | 2.3.1.54 | Acetyl-CoA + Formate <=> CoA + Pyruvate |

**S.3. PMD requires HRMS**

Once PMDs are calculated, linking these PMD_R_ to specific elemental compositions will provide valuable biological context. However, annotations of the elemental compositions of certain PMD are dependent on high resolution mass spectrometers, because low resolution instruments that only measure nominal mass may not be specific enough to distinguish elemental compositions. For example, a PMD 14 Da could be the addition or loss of a nitrogen atom or the addition of one oxygen atom and loss of two hydrogen atoms.

Here, we use HMDB ^4^ to demonstrate the effects of low resolution versus high resolution measurements in determining elemental compositions. PMD, as well as the elemental composition, were computed for the unique chemical formulas rounded to one, two, or three decimal places. As can be seen in Table S2, higher frequencies of a PMD are observed when rounding to less digits, suggesting the presence of false positives. As confirmation of the annotation accuracy, we determined how many of the PMDs in Table S2 resulted from a change in chemical formula linked with the appropriate PMD for the range of reported decimal places (Table S3). For example, of the 4381 ion pairs with a PMD of 14.016, > 98% of the pairs included an elemental change of +C2H. However, when two decimal places were reported, e.g. PMD of 14.02, only 63% of the 6875 ion pairs included an elemental change of +C2H. For the top 10 PMDs, accuracy > 95% was observed when the PMDs were rounded to three decimals, only ≥ 53% when rounded to two decimal places, and < 11% when only 1 or 0 decimals are used (see Table S3), confirming that high resolution mass spectrometry is required for qualitative PMD analysis and elemental composition annotation.

Table S2: Frequency of PMDs calculated from compounds in HMDB with decreasing mass accuracy.

| PMD  (digits = 3)^*^ | Frequency | PMD  (digits = 2) | Frequency | PMD  (digits = 1) | Frequency | PMD  (unit) | Frequency |
| --- | --- | --- | --- | --- | --- | --- | --- |
| 14.016 | 4934 | 14.02 | 8003 | 14.0 | 50419 | 14 | 156245 |
| 2.016 | 4909 | 2.02 | 7959 | 2.0 | 50467 | 2 | 156260 |
| 28.031 | 4878 | 28.03 | 7799 | 28.0 | 50797 | 28 | 155410 |
| 26.016 | 4229 | 26.02 | 7343 | 26.0 | 48517 | 26 | 154346 |
| 15.995 | 4214 | 15.99 | 7731 | 16.0 | 51278 | 16 | 155811 |
| 12.000 | 3861 | 12.00 | 7145 | 12.0 | 49335 | 12 | 155339 |
| 56.063 | 3861 | 56.06 | 6699 | 56.1 | 36417 | 56 | 151894 |
| 42.047 | 3771 | 42.05 | 6558 | 42.0 | 49808 | 42 | 153764 |
| 30.011 | 3698 | 30.01 | 6761 | 30.0 | 51241 | 30 | 154369 |
| 24.000 | 3689 | 24.00 | 6963 | 24.0 | 48099 | 24 | 154278 |

^*^ The top ten frequently occurring PMDs based on analysis of the data rounded to three decimal places were used as a reference. Frequency was then calculated for those PMDs when fewer decimal places were used.

Table S3: Effect of mass accuracy on elemental composition annotation accuracy^*^ of top ten selected PMDs from Table S2.

|  | PMD (digits = 3) | PMD (digits = 2) | PMD (digits = 1) | PMD (unit) |
| --- | --- | --- | --- | --- |
| +C2H | 0.98 | 0.63 | 0.10 | 0.03 |
| +2H | 0.97 | 0.62 | 0.10 | 0.03 |
| +2C4H | 0.99 | 0.61 | 0.09 | 0.03 |
| +2C2H | 0.98 | 0.58 | 0.09 | 0.03 |
| +O | 0.99 | 0.56 | 0.09 | 0.03 |
| +C | 0.99 | 0.56 | 0.08 | 0.03 |
| +4C8H | 0.97 | 0.56 | 0.10 | 0.02 |
| +3C6H | 0.98 | 0.58 | 0.08 | 0.03 |
| +C2HO | 0.95 | 0.54 | 0.07 | 0.02 |
| +2C | 0.99 | 0.53 | 0.08 | 0.02 |

^*^ The accuracy was calculated by dividing all of the frequency numbers of PMDs from Table S2 by the true numbers of compounds that contained the expected elemental composition. For example, 98% of the compounds with PMD 14.016 Da contain elemental compositions +C2H while only 63% of the HMDB compounds with PMD 14.02 Da contain elemental compositions +C2H. High resolution calculations of PMD show higher accuracy of elemental compositions.

Table S4. Demonstration of the selection of quantitative PMD pairs. Theoretical mass pairs [A, B], [C,D], and [E,F] are involved in the same PMD. Only [A, B] and [E, F] are considered static PMD and suitable for quantitative analysis since their intensity ratios were stable across sample 1 and sample 2.

|  | A^a^ | B | Intensity ratio | C | D | Intensity ratio | E | F | Intensity ratio |
| --- | --- | --- | --- | --- | --- | --- | --- | --- | --- |
| sample1 | 100 | 50 | 2:1 | 100 | 50 | 2:1 | 30 | 40 | 3:4 |
| sample2 | 1000 | 500 | 2:1 | 10 | 95 | 2:19 | 120 | 160 | 3:4 |

^a^ peak intensity of theoretical m/z.

**S.4 PMD network analysis**

PMD network analysis in this study describes a network connected by either sets of PMDs from known reactions or high frequency PMDs generated from the experimental data or databases. A local recursive search algorithm of PMDs to grow the network has been implemented in the pmd package. The identified peaks with specified PMDs are added to the network as secondary metabolites, and the process is repeated until all PMDs and extensions are exhausted. In addition, when the PMD network is generated from multiple samples, the paired masses’ intensities are required to have at least a moderate Pearson correlation coefficient (> 0.6 in this study) to build the linkage. To demonstrate this application, we obtained raw data from a published study to detect the biological metabolites of exposure to Tetrabromobisphenol A (TBBPA) in pumpkin ^14^. Those data were included in the enviGCMS package ^15^ (<http://yufree.github.io/enviGCMS/>, version 0.6.6) and re-analyzed to describe the PMD network analysis.

The pmd package ^10^ (<https://yufree.github.io/pmd/>, version 0.1.9) was developed with new features such as annotation using mass spectrometry detectable compounds from both KEGG and HMDB databases, quantitative PMD analysis for reactions, and PMD network analysis. It includes all of the PMD analysis described in this study.
